# Speciation history shapes patterns of assemblage species richness in birds

**DOI:** 10.1101/2024.08.28.610044

**Authors:** Bouwe R. Reijenga, Rampal S. Etienne, David J. Murrell, Alex L. Pigot

## Abstract

Speciation is the ultimate source of biodiversity. However, because most species arise in spatial isolation (allopatry), how speciation shapes patterns of co-occurring (sympatry) species richness remains unclear. Here we examine how the history of past speciation events influences the maximum sympatric species richness attained across passerine bird clades (n = 40 families). Using a phylogenetic model, we infer that the rate at which species assemble in sympatry is extremely slow, and on average millions of years after speciation onset. As a result, having accounted for clade size, sympatric species richness varies substantially across families, being highest in small or ancient clades comprised of older species that have had more time to accumulate in sympatry. While our analysis does not test the ecological or evolutionary processes causing the slow build-up of sympatric assemblages, our results show that speciation history has left an indelible legacy on current species richness patterns.

## Introduction

It has long been recognised that the number of species found at a particular place on the Earth’s surface reflects not only current environmental conditions but also the history of the environment and the lineages that occur there (Francis & Currie 2003; Harvey *et al*. 2020; Jetz *et al*. 2004; Jiménez-Alfaro *et al*. 2018; Sonne & Rahbek 2024). This effect of history on the richness of species assemblages operates over a range of spatial and temporal scales (Carrasco *et al*. 2022; Stephens *et al*. 2025). In some cases, the recency of major disturbance events allows the effect of history on community assembly to be observed directly, such as the recolonisation of plants and animals following the 1883 eruption of Krakatau (Bush & Whittaker 1991) or the retreat of northern hemisphere ice sheets at the end of the Pleistocene (Blois *et al*. 2010). Over deeper timescales, where community assembly must be inferred, phylogenetic approaches have been used to model the dynamics of species diversification and dispersal within and among biomes, regions and islands (Jetz & Fine 2012; Kozak & Wiens 2012; Miller *et al*. 2018; Quintero *et al*. 2023; Stephens & Wiens 2003; Swiston & Landis 2025; Valente *et al*. 2020; Wiens 2011). While such studies have advanced our understanding of what limits the accumulation of species within large or geographically isolated regions, they do not model how species assemble within the finer spatial grains (e.g., ∼25km to ∼100km grid cells) at which broad-scale gradients in species richness are typically mapped (Orme *et al*. 2005; Rahbek & Graves 2001). Understanding how history shapes the accumulation of species richness at these finer spatial grains requires consideration of both species origination and how these lineages come to assemble at a given location.

The origin of most species involves some form of spatial separation (allopatry or parapatry) that reduces gene flow between populations (Coyne & Orr 2004; Phillimore *et al*. 2008). As a result, cladogenesis becomes increasingly unlikely in geographic areas that are small relative to the spatial scale of dispersal (Coyne & Price 2000; Kisel & Barraclough 2010). In birds, for example, cladogenetic speciation is extremely rare or debatable in continuous areas smaller than Madagascar (1600 x 350 km) (Coyne & Price 2000; Phillimore *et al*. 2008). The richness of assemblages—here defined as the species co-occurring at a spatial grain below the limit of cladogenesis—is therefore generated through the colonisation of species from the surrounding regional pool and eroded via local extinction (MacArthur & Wilson 1967; Mittelbach & Schemske 2015; Pigot & Etienne 2015). While not directly adding species to assemblages, the history of speciation is still expected to shape patterns of species richness indirectly by structuring the species pool available for colonisation. First, differences in the cumulative number of speciation events across regions or clades, whether determined by differences in age (Kozak & Wiens 2012), rates of diversification or ecological limits (Rabosky 2012) will influence the size of the species pools from which local assemblages can be formed (Cornell & Lawton 1992). Second, and more rarely addressed, the timing of past speciation events will determine the age of the species in the pool and thus the time available to colonise local assemblages.

Previous studies analysing the transition of sister species to secondary sympatry (i.e. geographic range overlap) following speciation have shown that although there are exceptions, most sister species retain non-overlapping distributions for millions of years after the initiation of speciation (Alencar & Quental 2023; Anderson & Weir 2022; Pigot & Tobias 2015; Price 2010; Weir & Price 2011b). This lag in the attainment of secondary sympatry has been attributed to a variety of intrinsic and extrinsic mechanisms, including limited dispersal across geologically persistent geographic barriers, adaptation to different environments (Reijenga *et al*. 2023) and strong negative interactions among recently diverged lineages that have yet to acquire reproductive isolation or ecological compatibility (Pigot & Tobias 2015; Price 2010; Sexton *et al*. 2009; Weber & Strauss 2016). Regardless of the specific mechanism limiting successful colonisation, the time elapsed since speciation represents a potentially important constraint on the build-up of species within local assemblages. Specifically, where speciation events occurred deeper in time, lineages have had longer to overcome intrinsic and extrinsic barriers to sympatry, increasing assemblage richness (Stephens & Wiens 2003). Given the strong geographic and clade-level gradients in the temporal dynamics of speciation (Harvey *et al*. 2020; Jetz *et al*. 2012; Rabosky *et al*. 2018; Schluter & Pennell 2017), differences in the timing of past speciation events could leave an important imprint on present-day species richness. Yet, a previous focus on pairwise range overlap means that the effect of this ‘speciation legacy’ in limiting the species richness of entire assemblages has yet to be quantified.

A scenario in which ‘time for colonisation’ constrains assemblage species richness bears similarity to what has been termed the ‘time for speciation’ effect (Hutter *et al*. 2013; Stephens & Wiens 2003), but there are several important distinctions. First, according to the ‘time for speciation’ effect, clades which colonise a region earlier have more time to speciate and thus accumulate more species. Whereas the ‘time for speciation’ effect seeks to explain the size of the species pool, here our focus is instead on how the timing of speciation events within the species pool limits the assembly of local species assemblages (Wiens *et al*. 2011). Second, as an explanation for species richness patterns, the ‘time for speciation’ effect is generally regarded as an alternative to the ‘ecological limits’ model (Wiens 2011), in which species richness is regulated around an equilibrium set by competition between species for finite resources (Etienne & Haegeman 2012; Phillimore & Price 2008; Rabosky 2009). In contrast, here we argue that competition between closely related and ecologically similar species, provides one of the key mechanisms that preserves the imprint of speciation history on current patterns of assemblage richness.

Here we develop an approach to test the effect of speciation history in determining the build-up of species within local assemblages and, as a case study, apply this approach to predict differences in assemblage species richness across 40 family level clades of primarily North and South American passerine birds. By comparing assemblage species richness across different families, we exploit the fact that each of these clades has a unique history of speciation that should lead to predictable differences in sympatric diversity. Because the richness of local assemblages is constrained by the total number of species in each clade, and this effect of pool size has been widely studied (Kennedy *et al*. 2017, 2018; Kozak & Wiens 2012; Weir 2006), we focus on explaining the maximum proportion of species in each clade found in sympatry. The passerine radiations we use are ideal in this respect because they differ widely in both their phylogenetic branching patterns (i.e. speciation history), as well as the maximum proportion of species that co-occur—ranging from 12.8 to 83.3% of species for Passerellidae and the Onychorhynchidae respectively (Figure 1).

**Figure 1.**
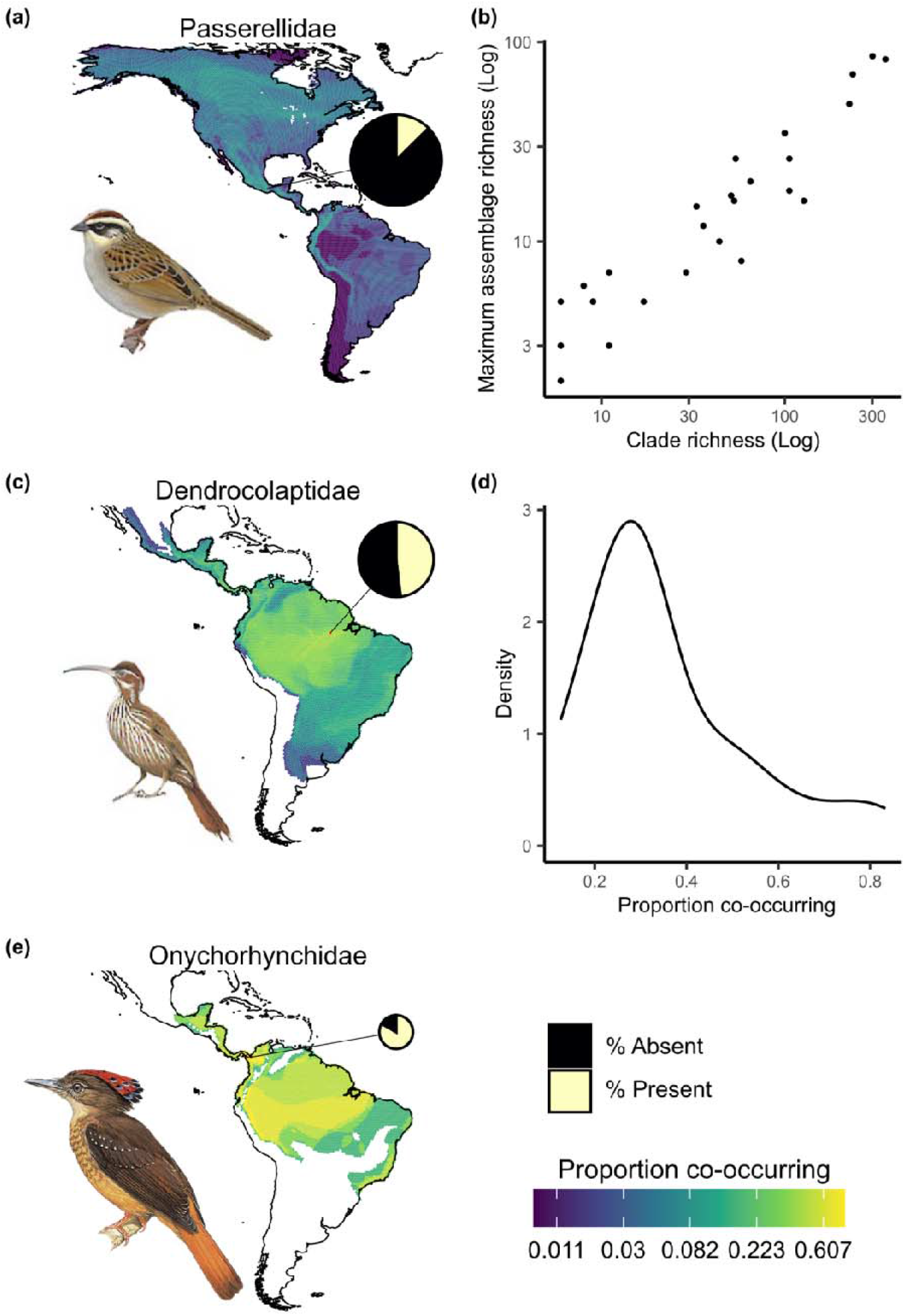
Species richness patterns of exemplary clades. (a, c, and d) assemblage species richness (as a proportion of clade richness) of three exemplar avian families showing low, intermediate, and high levels of co-occurrence. The size of the pie charts shows clade richness, and the black lines point towards the grid cell of highest richness. (b) the relationship between clade richness and maximum local species richness (*n*=40 families). An ordinary least squares regression estimates an intercept of -0.257 and slope of 0.761. (d) distribution of maximum (proportional) local species richness across clades.

**Figure 2.**
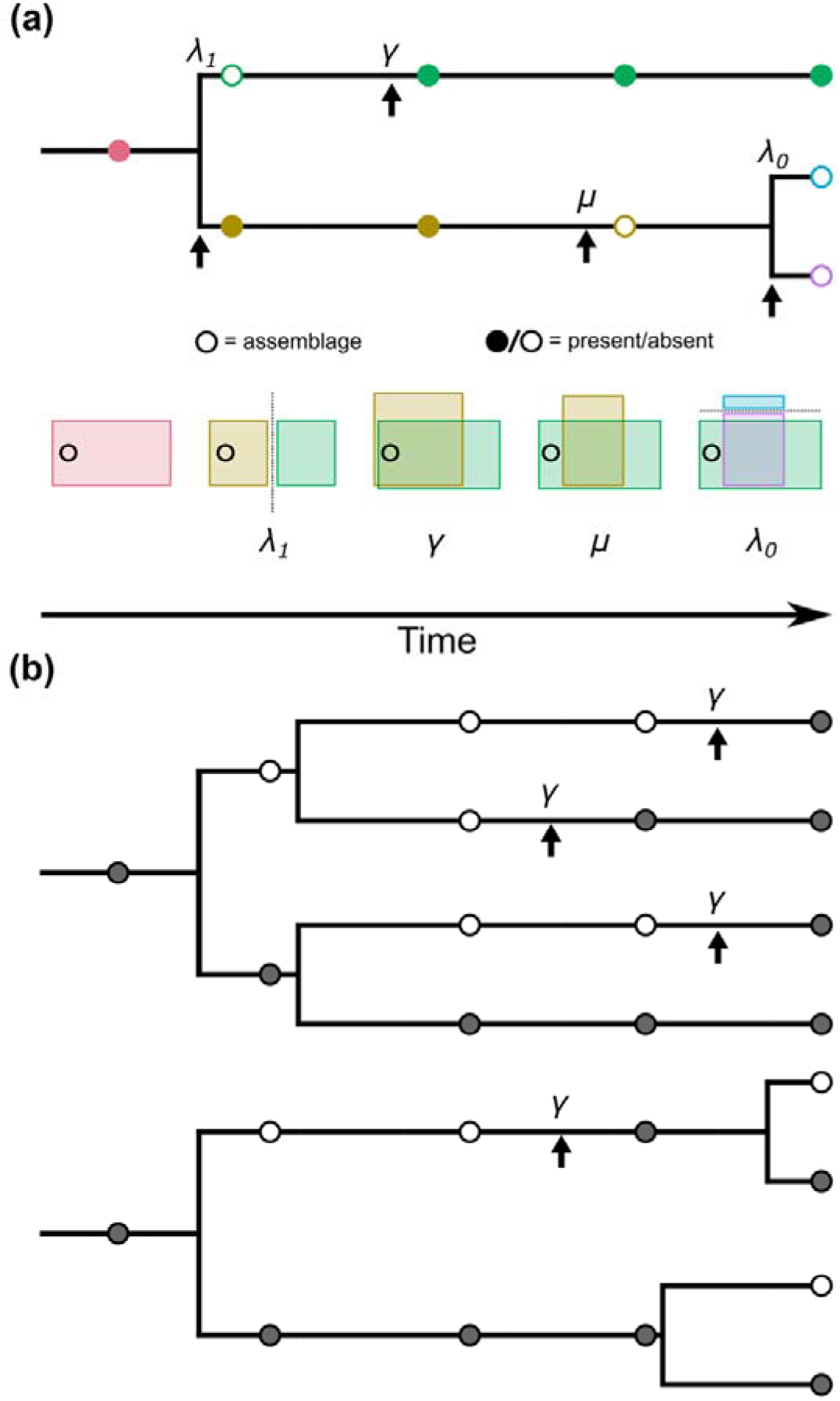
Visualisation of how speciation history influence assemblage richness. (a) conceptualisation of the link between allopatric speciation, range expansion and assemblage richness. Upward-pointing arrows show timing of events. When speciation occurs the ancestral geographic range (red rectangle) is divided into two (yellow and green), and at most one species will be present (λ_1_) in the local assemblage (circle). If the ancestor is not present, neither of the descendants (blue and purple) will be present (λ_0_) in the local assemblage. The build-up of local assemblage thus depends on the colonisation of species (γ), as well as local extinction (μ), as species undergo expansion (green) or contractions (yellow) of their geographic range. (b) two hypothetical clades, each with four species, with the top clade having older species on average than the bottom clade. Filled circles highlight the presence of a lineage in the local assemblage, and arrows indicate colonisation events. For both clade the ancestor was present in the assemblage, but the dynamics of allopatric speciation and colonisation result in a higher assemblage richness at present in the clade with older species.

Our analysis consists of the following key steps. First, for each family we use geographic range maps to characterise assemblages of sympatric species within hexagonal grid cells spanning ∼55km in width. This grid cell size corresponds to an area substantially smaller than the minimum scale of speciation in birds allowing us to assume that new species are added to assemblages exclusively through colonisation. We focus on the most speciose assemblage within each family, the location of which can, and often does, vary across clades. Second, given the set of species present in each of these focal assemblages and the speciation history captured by the phylogenetic branching patterns of each clade, we use a maximum likelihood-based framework to estimate the rates of colonisation γ and local extinction μ that best explain the present day composition of these assemblages. Our framework uses the Dynamic Assembly Model of Colonisation, Local Extinction and Speciation, ‘DAMOCLES’ (Pigot & Etienne 2015) that has previously been applied to infer the dynamics of community assembly within individuals clades, but here we extend this to model community assembly across multiple clades simultaneously. While DAMOCLES has previously been used as a null model for detecting non-stochastic community assembly processes (e.g. environmental filtering or competition)(Marx *et al*. 2017; Pigot & Etienne 2015; Pinto-Ledezma *et al*. 2019), here we instead apply this approach to infer γ and μ, and are agnostic to the ecological processes that regulate these dynamics. We note that without a complete fossil record for each assemblage, the timepoint at which each species colonised these locations is unknown. However, given the large spatial scale of speciation relative to our assemblages, we can be confident that following each speciation event, at least one of the daughter species must initially have been locally absent. DAMOCLES uses this insight to constrain when colonisation could have occurred and to estimate the rate that best explains current assemblage structure. Third, we use our empirically-derived estimates for γ and μ to stochastically simulate colonisation and local extinction events over the history of each clade and generate expectations for the current local assemblage richness attained by each family. In our main model we constrain γ (and μ) to be identical across all clades, so that any differences in expected assemblage richness are due solely to differences in the speciation history of each clade. In this way, we test how well speciation history can predict observed variation in assemblage species richness across clades, as well as how assemblage species richness varies with a suite of phylogenetic metrics describing the historical dynamics of speciation. Throughout, we compare the fit and predictive ability of this model, assuming a single universal rate of γ (and μ), to a model where these parameters can vary independently across families, to test how relaxing our assumption of equivalent rates alters our conclusions. Finally, we compare this model to a scenario where γ is fixed at a very large value and only μ is estimated. In this counterfactual scenario, the colonisation of species assemblages occurs almost instantaneously relative to the timescale between speciation events and so speciation history has no effect on current assemblage composition or richness.

## Materials and Methods

### Phylogenetic and geographic range data

We compiled a dataset consisting of oscine and suboscine passerines predominantly endemic to the Americas. These clades were chosen because their high species richness (n = 2081 species) provides sufficient statistical power, and because phylogenetic and geographic data is near complete. Evolutionary relationships and divergence times were obtained from two time-calibrated phylogenies containing >95% (oscines) and >98% (suboscines) of described extant species (Barker *et al*. 2015; Harvey *et al*. 2020). To examine how clade-specific speciation histories impact present-day assemblage richness the phylogenies were subdivided into family-level clades with at least five species, resulting in 7 oscine and 18 suboscine clades. This balanced sufficient variation in patterns of assemblage richness with large enough clades to reliably calculate phylogenetic metrics.

To identify the species co-occurring in each family we used expert delineated range maps (BirdLife International and Handbook of the Birds of the World 2020). Compared to field inventories which are often incomplete, range maps provide a comprehensive characterisation of species richness patterns albeit at a coarser spatial resolution. Because the phylogenies and range maps were based on different taxonomies, we aligned them by using the phylogenetic taxonomy and merged or split species geographic ranges accordingly. Range maps were then projected onto an equal area hexagonal grid (∼2600 km^2^ per cell), where each hexagon has a side length of ∼31.6 km and the distance between cell centroids is ∼55km (Barnes & Sahr 2017). This grain size is comparable to those used in macroecological analyses (Jetz & Fine 2012; Pigot *et al*. 2016; Rahbek 2005), and is much smaller than the minimum area required for *in situ* speciation to occur in birds (Coyne & Price 2000; Kisel & Barraclough 2010), thus ensuring that co-occurring species arrived through colonisation. For each family-level clade, we identified the grid cell of maximum species richness and recorded its composition. For the small number of species lacking expert range maps (n = 5, equally distributed among clades of >53 species each), we used auxiliary information (e.g., occurrence records) to confirm their presence/absence in the grid cell of maximum richness for each family. While acknowledging that expert range maps are subject to errors of commission at small grain sizes (Hurlbert & Jetz 2007), we repeated our analyses using a finer grid (∼96 km^2^ area with cell centroids ∼10 km apart and sides of ∼6 km) that reduces the chance that allopatric species separated by narrow geographic barriers are scored as co-occurring.

### Estimating the dynamics of community assembly

To test if speciation legacies can explain differences in assemblage richness across clades we applied and modified the DAMOCLES framework (Pigot & Etienne 2015). DAMOCLES models the assembly of a single community from a dynamically evolving regional pool via the processes of colonisation (γ) and local extinction (μ) (Figure 1), assuming constant rates through time and among lineages. Speciation is not modelled. Instead, the composition of the regional pool at time *t* reflects the lineages in an empirical reconstructed phylogeny that are extant at time *t*. DAMOCLES assumes that cladogenesis will result in one daughter lineage being present and one absent if the parent lineage is present in the assemblage, and that both will be absent if the parent is absent (Figure 1b). Given this model, the phylogeny and a vector of the presence (1) and absence (0) of each species in the grid cell of maximum richness, we estimated γ and μ using maximum-likelihood.

Obtaining the likelihood of an assemblage (i.e. the vector of presence/absence) resembles the Felsenstein pruning algorithm (Felsenstein 1981), detailed in Pigot & Etienne (2015). Operating from extant tips to the root, we track each lineage back in time, conditioning on observed presence (1) and absence (0) at the tips. We define *p*_*if*_ (*t*) as the probability that the descendants of a lineage are in state *f* at the present (time 0), given that the lineage was in state *i* at time *t* in the past. The change in these probabilities along a branch is described by a system of Ordinary Differential Equations (ODEs), which are integrated backward in time until a node is reached. At each node, probabilities of the two descendant lineages are combined, and the two lineages are pruned. This process is repeated until there are no more branches left to prune. The likelihood at the root is calculated by combining *p*_*0f*_ and *p*_*1f*_, with the combination determined by the prior distribution on the root state. Assuming complete ignorance, we compute the likelihood as the sum of these probabilities. The values of γ and μ that best explain a community are obtained by maximising this likelihood. To ensure robust inference, we fitted models by using multiple solvers to confirm numerical stability, using multiple starting parameter values and multiple optimisation algorithms to ensure convergence to the global likelihood maximum.

### Testing the effect of speciation history on assemblage richness

We tested the effect of speciation history on maximum assemblage richness across clades via three model scenarios. First, a ‘historical global rate’ model estimated a single γ and μ across all clades, such that differences in (proportional) assemblage richness across clades arise entirely from differences in speciation history and time for colonisation. Second, a ‘historical variable rate’ model estimated γ and μ separately for each family, thus relaxing the assumption that these dynamics are identical across clades. For both scenarios, the likelihood was conditioned on assemblages containing at least one species. Finally, we evaluated a ‘non-historical’ null model only estimating μ and fixing γ at a very high rate (γ = 1000), representing effectively no lag time to colonisation after speciation. Failure to reject this null model would indicate no detectable effect of speciation history on the richness of species assemblages. We compared the fit of the ‘historical global rate’, ‘historical variable rate’ and ‘non-historical’ model using AIC. Parametric bootstrapping was used to estimate the 95% CI for the ‘historical global rate’ and ‘historical variable rate’ model parameter estimates (Table S1).

We assessed model adequacy of the ‘historical global rate’, ‘historical variable rate’ and ‘non-historical’ scenarios by comparing their ability to reproduce the observed assemblage richness of each family and also to predict how assemblage richness varies according to different phylogenetic metrics describing the history speciation within each clade. For each model, we simulated lineage colonisation and local extinction of assemblages for each family using estimated γ and μ values. Simulations were implemented via a Gillespie algorithm (Gillespie 1977), and modelled from the root of the family phylogeny at *t* = 0 until the present. Lineages transitioned from being absent (0) to being present (1) via colonisation (0 ⍰ 1) and local extinction (1 ⍰ 0) events, assigning the root state with equal probability. Waiting times between events (δ) were drawn from an exponential distribution based on the summed per-lineage rates. At time *t* + δ, the event type and undergoing lineage was drawn based on the individual rates. This framework was already incorporated in DAMOCLES, but we re-wrote this in *Rcpp* (Eddelbuettel & Balamuta 2018). Simulations were repeated 2500 times per clade to obtain the mean and 95% confidence interval (95% CI) in simulated assemblage richness under the ‘historical global rate’, ‘historical variable rate’ and ‘non-historical’ scenarios.

We used the 95% CI in simulated assemblage richness to identify families with significantly higher or lower assemblage richness than expected under each model. We then tested how both the observed and mean simulated assemblage richness varied across clades according to the following phylogenetic metrics. For each family we quantified: (*i*) the crown age (Myr), (*ii*) tree imbalance quantified using Colless’ index standardised for tree size and richness, where higher values indicate greater imbalance (Bortolussi *et al*. 2006; Mooers & Heard 1997), (*iii*) diversification (speciation – extinction) rate change (ρ), which measures slowdowns (ρ < 0) or speed-ups (ρ > 0) in diversification through time (Etienne & Rosindell 2012; Janzen & Etienne 2024; Pigot *et al*. 2010), and (*iv*) mean terminal branch length (mbl) as a measure of the average age of extant species. Each metric was tested as a predictor of proportional assemblage richness across families using quasibinomial GLMS with a logit link (Douma & Weedon 2019). We fit separate models for the observed assemblage richness and the mean simulated richness for each of our three scenarios (i.e. ‘historical global rate’, ‘historical variable rate’ and ‘non-historical’). Analysis was restricted to clades with more than five species, because calculating phylogenetic properties for very small trees becomes less informative.

## Results

### The dynamics of community assembly

According to the ‘historical global rate’ model, the maximum likelihood per lineage rate of colonisation and local extinction is γ = 0.120 Myr^-1^ and μ = 0.178 Myr^-1^ respectively. This equates to a mean lag time from speciation to the colonisation of the focal assemblage of 8.3 Myr. A similar dynamic is inferred using the ‘historical variable rate’ model. In the ‘historical variable rate’ model, estimates of γ (0.04 to 100 Myr^-1^) and μ (0 to 187 Myr^-1^) vary substantially across different families (Table S1). However, families with very high estimated rates of γ and μ are small and thus have low information content such that rate estimates are highly uncertain (Table S1). As a result, the mean lag-time to colonisation weighted for clade richness under the ‘historical variable rate’ model (7.9 Myr) is similar to that of the ‘historical global rate’ model (8.3 Myr). Comparison based on AIC indicates that the ‘historical global rate’ model (AIC_*γμ*_ = 2333.398), assuming a single γ and μ across all clades (n = 2 parameters), is better supported than the more parameter-rich ‘historical variable rate’ model (AIC_*V*_ = 2350.29, n = 50 parameters, corresponding to separate γ and μ for each clade). Importantly, both the ‘historical global rate’ (AIC_*γμ*_ = 2333.398) and ‘historical variable rate’ model (AIC_*V*_ = 2350.29) provide a substantially better fit to the data than a ‘non-historical’ model in which rapid colonisation and local extinction dynamics erase the effects of speciation history (AIC_*NH*_ = 2379.482). While exact rate estimates vary, these conclusions were consistent when using a finer grain size to define species assemblages (96 km^2^, Figure S3), and when selecting focal assemblages matching the mean level of richness observed within each clade rather than the maximum species richness (Figure S5).

### The effect of speciation history on assemblage richness and phylogenetic metrics

Posterior simulations further supported the ‘historical’ models, with observed proportional assemblage richness falling within the 95% CI for 21 (‘historical global rate’) and 25 (‘historical variable rate’) of 25 clades. In contrast, the ‘non-historical’ model reliably predicted assemblage richness in only 16 out of 25 clades (Figure 3c). Across clades, the maximum proportion of species that co-occur increases with clade age and mean terminal branch length, and declines with clade richness (Figure 4, Figure S4, Figure S6). There was no effect of phylogenetic imbalance nor family-level ρ on the maximum proportion of co-occurring species. These patterns were also predicted when we simulated assemblages under both the ‘historical global rate’ and ‘historical variable rate’ models. In contrast, the ‘non-historical’ null model predicts little variation in proportional assemblage richness (Figure 3c) and no correlation with phylogenetic metrics (Figure 4). Using assemblages of mean rather than maximum species reduces the variation in richness observed across clades, but does not qualitatively alter our conclusions (Figure S6).

**Figure 3.**
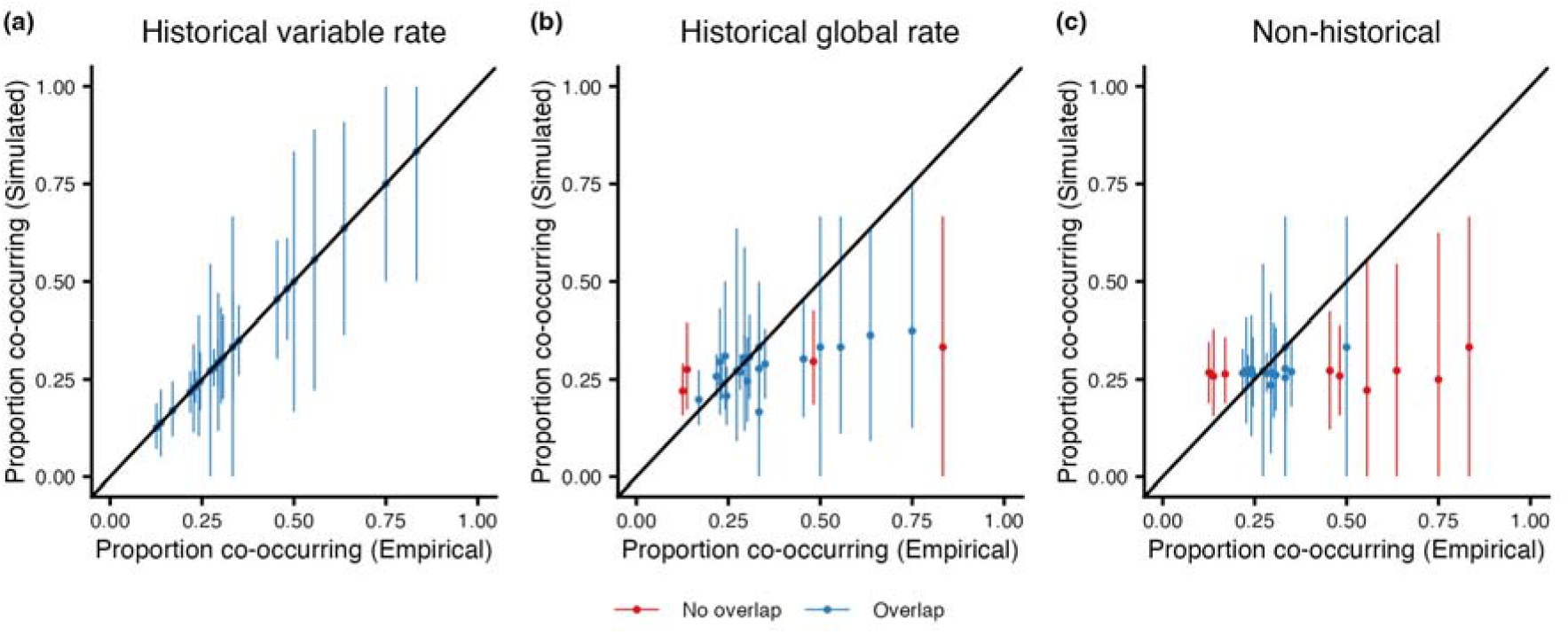
Observed variation in assemblage richness of passerine clades compared to model predictions. Assemblage richness is plotted as a % of the total number of species in each clade. Model predictions are shown for (a) the ‘historical variable rate’, (b) ‘historical global rate’, and (c) the ‘non-historical’ model. The 1:1 line shows where the empirical the predicted richness matches. Bars represent the 95% confidence intervals of predicted assemblage richness from 2500 replicate simulations of each model. Colours indicate if empirical assemblages fall within (blue) or outside (red) of model expectations.

**Figure 4.**
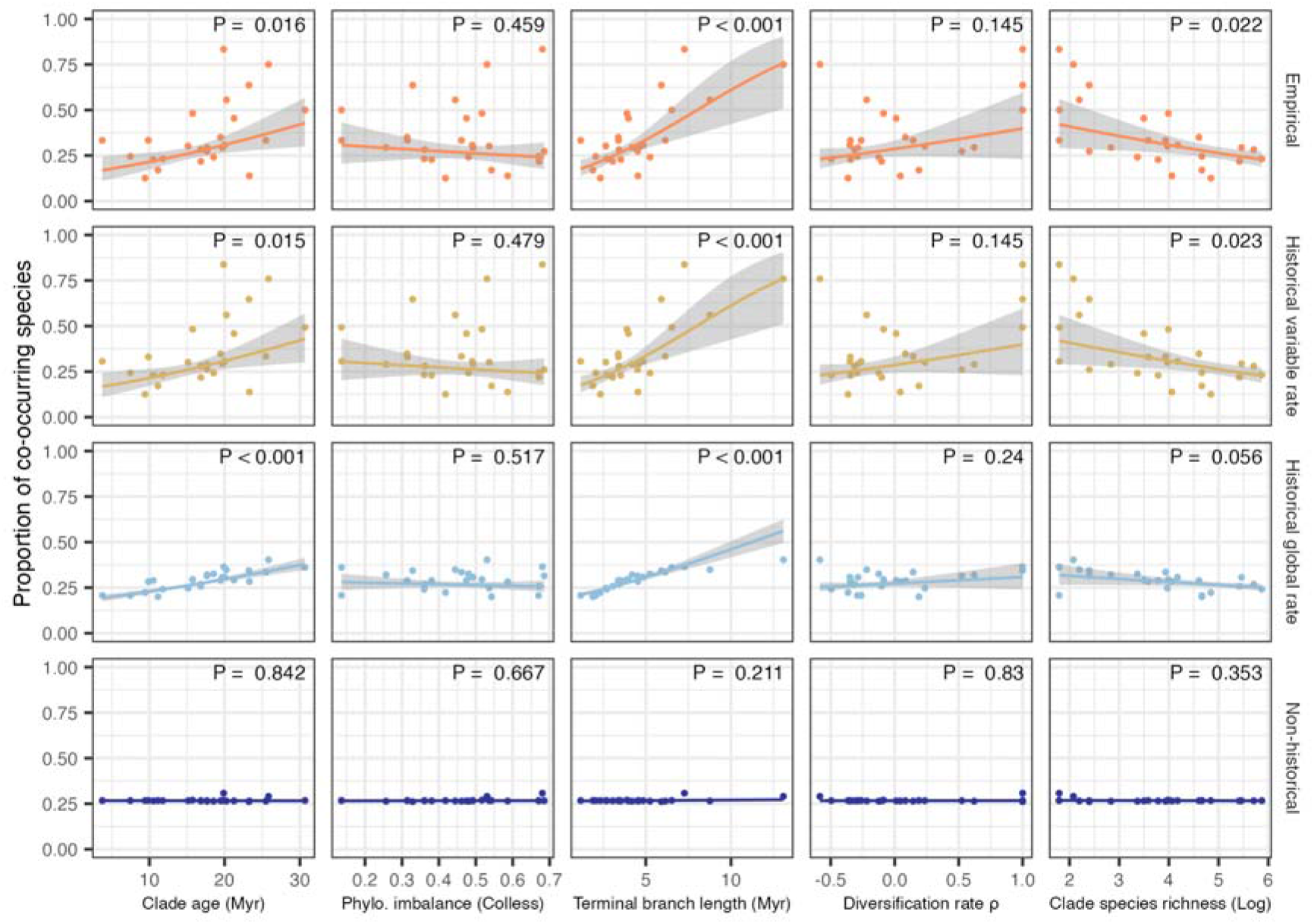
Relationship between phylogenetic metrics and assemblage species richness for real bird clades and model simulations. Colours and rows denote the empirical relationships for all clades (orange) and relationships simulated under the ‘historical variable rate’ (yellow), ‘historical global rate’ (blue) colonisation rate and ‘non-historical’ (dark blue) model. For the simulation scenarios the mean proportion of co-occurring species across 2500 replicated simulations are plotted. Columns represent the five phylogenetic metrics capturing different aspects of clade evolutionary history: The crown age of the clade, phylogenetic imbalance (Colless’ index), mean branch length of the extant species, ρ a measure of temporal change in diversification rates through time where positive (negative) values indicate an increase (decrease), and log-transformed clade species richness. Shaded areas represent 95% confidence intervals of fitted generalised linear models with quasibinomial error distribution and logit link function weighted for clade richness.

## Discussion

The evolutionary history of speciation, extinction and colonisation has been argued to be of fundamental importance in explaining the current richness of species assemblages (Mittelbach & Schemske 2015; Ricklefs 1987). Here, by expanding upon a dynamic model of community assembly that incorporates these processes, we show that the history of past speciation events has left a clear legacy in the number of species co-occurring within assemblages of passerine birds.

Our model incorporates the historical legacy of speciation by assuming that upon speciation daughter lineages do not co-occur and that the build-up of species richness in sympatry thus depends on colonisation, an assumption that is widely supported across animals and plants (Coyne & Orr 2004; Coyne & Price 2000; Olivares *et al*. 2025; Phillimore *et al*. 2008). Focussing on the most species rich assemblage within each bird family, we estimate that the mean lag-time (i.e., 1/γ) between the speciation of a lineage and its incorporation into this focal assemblage is approximately 8 Myr. This suggests that build-up of species in sympatry occurs over timescales comparable to macroevolutionary rates of speciation. For instance, the average per-lineage waiting time between speciation events leading to extant birds is 6.25 Myr (i.e. 1/0.16 species Myr^-1^)(Jetz *et al*. 2012). While the stochastic nature of colonisation will result in some species attaining sympatry rapidly, the slow average rate of colonisation suggests that the timing of past speciation events strongly constrains the build-up of species richness within assemblages. Indeed, we show that differences in the history of speciation across clades leads to predictable variation in assemblage richness. In particular, a higher proportion of species co-occur in older clades as these contain older species that have had more time to accumulate in sympatry. In contrast, while larger clades have more speciose assemblages in absolute terms, the proportion of species that co-occur is reduced. This is because, large clades will tend to be those that have experienced many recent speciation events, leading to younger species that have had less time to accumulate in sympatry. Accordingly, a non-historical null model in which local species temporal turnover is rapid, and that leads to assemblages with random phylogenetic structures, is rejected.

Our results show that patterns of assemblage richness in birds are best explained by a model in which the build-up of species in sympatry is a slow process, but this finding begs the question of what limits the rate of colonisation. One possibility is that species adapt to spatially non-overlapping habitats and this prevents them from occupying the same location in the landscape (Marx *et al*. 2017). Such environmental filtering is indeed expected across the families of passerine birds in our analysis (Pinto-Ledezma *et al*. 2019), but is unlikely to explain our results. Simulations show that widespread environmental filtering causes both young and old species to be absent from local assemblages. This absence of a relationship between species age and the probability of being in sympatry would lead to our models, which assume equal rates of colonisation (and local extinction) across lineages, inferring rapid local turnover, rather than the slow rates of colonisation that we estimate (Pigot & Etienne 2015).

A more likely explanation for why the legacy of speciation persists in present day assemblages is that range expansions leading to secondary sympatry are strongly constrained by a combination of dispersal limitation and biotic interactions (Tobias *et al*. 2020). While birds are relatively dispersive organisms that can expand their ranges rapidly across continuous landscapes, geographic barriers that initiate speciation but then persist in the landscape for millions of years (e.g. mountain ranges, rivers) are likely to continue to enforce spatial isolation long after speciation is completed (Naka & Brumfield 2018; Pigot & Tobias 2013). For example, numerous sister species of Neotropical birds are isolated on either side of the Andes, on separate mountain peaks and ridges or by wide habitat breaks (e.g. Amazonian versus Atlantic forests) (Mikkelsen *et al*. 2025; Pérez-Emán 2005; Weir & Price 2011a). Beyond dispersal limitation, competition for ecological resources and reproductive interference are key constraints in the transition to secondary sympatry in birds (Anderson & Weir 2021; Pigot *et al*. 2018; Weir & Price 2011b). Complete reproductive isolation typically evolves over millions of years in birds (Price 2008; Price & Bouvier 2002). Secondary contact between incipient species before reproductive isolation has completed can either cause the formation of hybrid zones that maintain abutting (i.e. parapatric) ranges or fusion back into a single lineage (Cooney *et al*. 2017). In ovenbirds and woodcreepers, two of the families we analyse here, transition rates to sympatry are faster among sister species that have rapidly diverged in ecomorphological traits linked to resource use (e.g. beak size) and where competition is presumably weak (Pigot & Tobias 2013). However, most sister species in these clades have highly conserved morphologies resulting in estimates of waiting times to secondary sympatry that are significantly longer than the age of most species, equating to essentially indefinite spatial exclusion (Pigot & Tobias 2013). Limited dispersal, incomplete reproductive isolation and competition exclusion are not mutually exclusive explanations and are likely to interact to explain the very slow build-up of species in sympatric assemblages and the persistent signature of speciation history reported here.

If negative biotic interactions between species are important in delaying the colonisation of assemblages over macro-evolutionary timescales, then the question becomes: is the richness of local assemblages unbounded, increasing over time as species escape competitive constraints by expanding into novel regions of ecological niche space (Harmon & Harrison 2015)? Or, are ecological opportunities bounded, with assemblage species richness regulated around a dynamic equilibrium (Rabosky & Hurlbert 2015)? In the unbounded non-equilibrium scenario, slow rates for colonisation can legitimately be interpreted as a constraint in the build-up of species in sympatry, albeit one ultimately controlled by ecological interactions. By contrast, in the bounded scenario, rates of colonisation could have been rapid in the past and only slowed down recently as local niche space has become filled (Rabosky & Hurlbert 2015). In this case, it would be incorrect to view the positive effect of species age on community membership that we observe across birds, as evidence of a long lag-time from speciation to colonisation as our constant rate models assume. Instead, the positive relationship between species age and community membership would be due a priority effect, whereby species that arose earlier, and were able to colonise assemblages first, have pre-empted niche space preventing younger species from colonising (Reijenga *et al*. 2021; Stroud *et al*. 2024).

Given these different interpretations for the cause of speciation legacies on assemblage richness, resolving whether local assemblage richness is bounded or unbounded, is a critical next step. This will require moving beyond the constant rate models used here to develop models that explicitly consider how rates of colonisation and/or local extinction vary over time or according to current levels diversity and niche structure (Valente *et al*. 2015). Nevertheless, we contend that a bounded scenario is unlikely to account for the patterns we demonstrate. Priority effects are expected to operate most strongly among closely related lineages that occupy similar niches within the same assemblage (Fukami 2015). It is therefore hard to envisage how priority effects could drive the relationship between assemblage species richness and phylogenetic structure that we observe across avian families, and why families with on average younger species have proportionally fewer species in sympatry. Slow rates of assemblage colonisation driven by a combination of habitat filtering, dispersal and biotic interactions would appear a more parsimonious explanation for the effect of speciation history observed both within and across clades.

While ultimately requiring further analysis with models that explicitly incorporate biotic interactions, our results suggest that patterns previously put forward as evidence for ecological limits to assemblage species richness could simply be explained by speciation history. For example, Weir (2006) showed that clades of Amazonian birds showing stronger slowdowns in diversification rate over time measured by the γ-statistic (Pybus & Harvey 2000) contained more speciose assemblages, and interpreted this as evidence that the saturation of niche space within local assemblages was inhibiting ongoing range expansions required for speciation (Weir 2006). However, the γ-statistic is dependent on clade size and, as we have shown here, the proportion of species expected to co-occur declines with clade richness even in the absence of diversity-dependence if there is a long lag-time between speciation and assemblage colonisation (Figure S8). Indeed, using an alternative metric that is independent of clade size (ρ) (Pigot *et al*. 2010), we find no relationship between clade slowdowns and the proportion of species that co-occur locally (Figure 4). While our results do not rule out the possibility that ecological limits have caused a slowdown in species diversification rate (Rabosky & Hurlbert 2015), they show that patterns in assemblage species richness should be evaluated against models incorporating the geographic context of speciation before being interpreted as evidence for or against ecological limits.

Another way in which the effects of ecological limits on species assemblages have been inferred is to consider the Lineage-through-time (LTT) plot of the ‘community phylogeny’ which tracks how the number of phylogenetic branches representing the species present in an assemblage has accumulated over time (Price *et al*. 2014). Just as LTT plots have been used to detect slowdowns in species diversification within clades (Etienne *et al*. 2012; Etienne & Haegeman 2012), it has been suggested that slowdowns in the LTT plot of a community phylogeny could indicate a declining rate of colonisation over time due to the filling of local niche space (Price *et al*. 2014; Weir 2006). However, we suspect that such a pattern could also arise from the historical legacy of speciation. Specifically, if the rate at which species colonise local assemblages following speciation is slow but constant, relatively few closely related lineages will co-occur leading to the appearance of a slowdown.

To test this possibility, we performed a post-hoc analysis, pruning the phylogeny of each family to the species present in the focal assemblage. We repeated this for the replicate assemblages simulated under our three models (the historical global rate, historical variable rate and non-historical null model), all of which assume constant rates of colonisation over time. We identified families exhibiting stronger or weaker slowdowns in the community phylogeny than expected under each model using both a rank envelope test (Mrkvička *et al*. 2017; Murrell 2018) and the 95% confidence interval in expected ρ (see Supplementary materials). A stronger slowdown would provide evidence that rates of colonisation have declined towards the present, as would be expected under the ecological limits model. We found that across avian families community phylogenies consistently show temporal slowdowns in branching rate (Figure 5, Figure S7). However, in 92% of clades (except Calyptomenidae and Tyrannidae), these slowdowns are no different from that expected under the ‘non-historical’ null model. Because the ‘non-historical’ null model generates assemblages that are random with respect to phylogenetic structure, this suggests that slowdowns arise simply from pruning the phylogenetic tree to only contain sympatric species (Figure S7). This artefact is identical to the effect of incomplete taxonomic sampling in causing the appearance of slowdowns in species diversification rate (Cusimano & Renner 2010). When the observed patterns are compared to the ‘historical global rate’ and ‘historical variable rate’ model, no clades or only a single clade (Calyptomenidae) show stronger slowdowns than expected respectively, and some clades (Emberizidae, Parulidae, and Thraupidae) show weaker slowdowns than expected. Thus, a combination of (1) pruning the phylogeny to co-occurring lineages and (2) the lag time between speciation and colonisation of assemblages is sufficient to explain almost all the slowdowns in community LTT plots without invoking ecological limits. While it has long been recognised that slowdowns in diversification rates could arise because speciation takes time (i.e. protracted speciation (Etienne & Rosindell 2012)), here we show that slowdowns in community phylogenies can arise because speciation generates spatially isolated lineages and colonisation of the same communities takes time.

**Figure 5.**
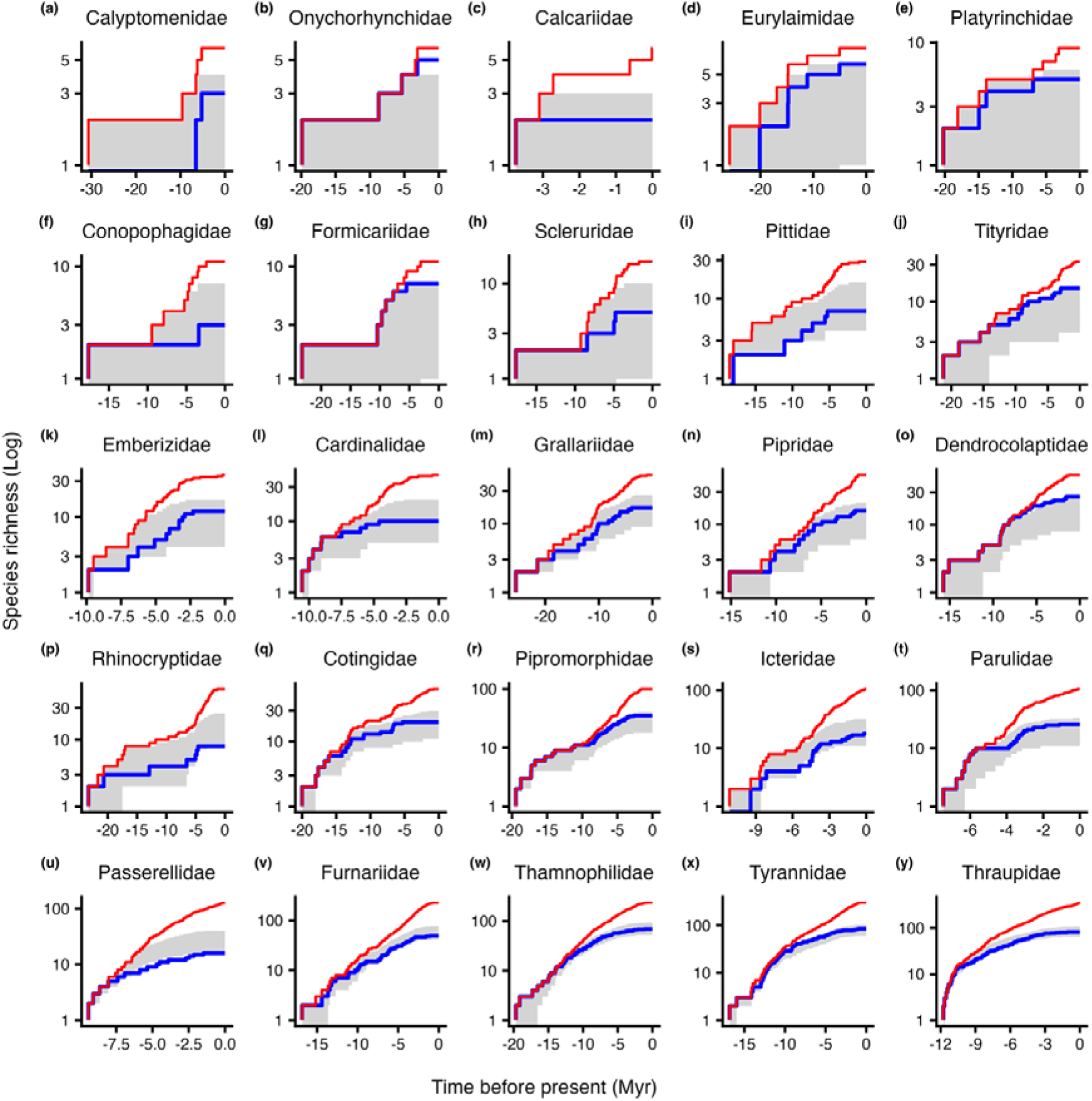
Clade and assemblage lineage-through-time (LTT) plots showing the accumulation of lineages through time within each family (red) and assemblage (blue, i.e., grid cell of maximum richness) compared to model predictions. Shaded areas correspond to the 95% confidence intervals of a rank envelope (Mrkvička *et al*. 2017) for the assemblage LTT expected under the ‘historical global rate’ model (*n* = 2500 replicate simulations). Families are ordered from smallest (Calyptomenidae, 6 species) to largest (Thraupidae, 354 species).

Our analysis shows that speciation history leads to predictable variation in assemblage richness across clades, but our study is subject to several caveats. We analysed patterns of sympatric species richness at grain sizes consistent with those used in broad-scale analyses of species richness gradients, but that are substantially larger than in many studies of community structure. As the area over which assemblages are defined is narrowed, we expect rates of local extinction and temporal turnover to increase (Dornelas *et al*. 2014; MacArthur & Wilson 1967), leading to a weaker imprint of speciation history in determining assemblage richness. For example, the application of DAMOCLES to plant assemblages on small Alpine summits suggest either rapid species turnover or substantial environmental filtering dominates assemblage structure (Marx *et al*. 2017). Even at the spatial scale we use, much of the variation in levels of sympatry observed across bird families remains unexplained. Given the simplistic nature of our models, this is not surprising and likely reflects differences across clades in factors regulating community assembly, such as the strength of environmental filtering, dispersal ability or negative biotic interactions. For example, the maximum sympatric species richness (*n* = 26) obtained by woodcreepers (Dendrocolaptidae) and (*n* = 5) Royal flycatchers (Onychorhynchidae) is significantly greater than expected given the phylogenetic branching patterns and ages of species in this clade. In contrast, the maximum sympatric species richness of New World Sparrows (Passerellidae) and Tapaculos (Rhinocryptidae) is significantly lower than expected given the phylogenetic branching patterns and ages of species in these clades (Figure 3b). By identifying clades where levels of sympatry depart from that expected due to speciation history alone, our models could help identify the additional factors limiting species richness. Future studies could also advance beyond this by explicitly modelling how rates of colonisation and local extinction vary both across lineages and geographic space within clades depending on the local environment (e.g., ecosystem productivity)(Pigot *et al*. 2016), the dynamics of different geographic barriers (e.g. ice sheets versus rivers)(Smith *et al*. 2014; Weir & Schluter 2004), species traits (e.g., dispersal ability (Sheard *et al*. 2020) and arrival order (Fukami 2015; Reijenga *et al*. 2021). While our parsimonious model shows that speciation history has left an indelible signature on current patterns of assemblage richness, fully understanding the role of evolutionary history in community assembly will require more complex models incorporating interspecific interactions both within and between clades (Reijenga *et al*. 2023), and the potential feedbacks between local assemblages and the dynamics of diversification and niche evolution within the species pool (Rabosky 2013; Reijenga *et al*. 2021; Storch & Okie 2019).

## Acknowledgements

We thank James Rosindell and Luis Valente for comments and discussion on an earlier version of this work.

