## Supplementary information for "Speciation history shapes patterns of assemblage species richness in birds"

*Lineage-through-time post-hoc analysis*

Lineage-through-time (LTT) plots track branch accumulation in a phylogenetic tree through time. Deviations in LTT shape from a constant-rate species diversification model could indicate non-random processes. For example, a slowdown in branch accumulation has been interpreted as a potential signal of adaptive radiation, where an initial burst of speciation is followed by a slowdown in diversification as ecological niches are filled (Etienne *et al.* 2012; Etienne & Haegeman 2012). Similar logic has been applied to LTT plots constructed from only sympatric lineages in an assemblage (Price *et al.* 2014; Weir 2006), what we here term ‘community phylogenies’. A slowdown in the LTT plot of a community phylogeny could indicate declining colonisation over time due to the local filling of niche space. However, similar patterns could be expected from speciation history alone: if colonisation following speciation is slow but constant, few closely related lineages will co-occur. To test this, we pruned each family phylogeny to include only the species present in the grid cell of maximum richness (i.e. a community phylogeny) and constructed the LTT plot for these retained lineages. We compared the shape of these LTT plots to those expected under our constant rate models. Specifically, we used the empirically-derived estimates of γ and μ obtained by fitting our three DAMOCLES scenarios to the observed data (‘historical global rate’, ‘historical variable rate’, and ‘non-historical’ model) to stochastically simulate colonisation and local extinction events over the history of each clade and to generate current local assemblages. We conducted 2500 replicate simulations for each model and clade. For each repetition, we pruned the family phylogeny to retain only those species currently present in the simulated local assemblages, this mimicking the same procedure as was used for the observed data. For the LTT plots simulated under each model, we constructed 95% confidence intervals via a rank envelope to deal with statistical non-independence (Mrkvička *et al.* 2017; Murrell 2018). Additionally, we calculated ρ for empirical and simulated community phylogenies and generated 95% confidence intervals in the expected values of ρ for each model against which the empirical values could be compared.

Significant departures of the empirical LTT plots and values of ρ from those expected would indicate that the observed patterns cannot be explained by a scenario in which colonisation (and local extinction) rates are constant over time. The ‘non-historical’ model assumes high rates of species temporal turnover and produces assemblages that are random with respect to phylogeny (Pigot & Etienne 2015). This is thus equivalent to randomly sampling species from the phylogeny. Failure to reject the ‘non-historical’ model would therefore indicate that a slowdown in the empirical LTT plot is simply the result of sampling a subset of species in the phylogeny. This effect has previously been noted with respect to detecting spurious slowdowns in diversification rates when not all species in the clade are sampled and thus included in the phylogenetic tree. This is because missing species results in the pruning of recent branches in the phylogeny while deeper branches tend to be retained (Cusimano & Renner 2010). A stronger slowdown that expected under the ‘non-historical’ model would indicate that processes beyond subsampling the phylogeny are contributing to LTT plot shape. To determine whether this is due to a slowdown in colonisation rate or is simply a legacy of speciation we can compare the LTT plot shape to that expected under the ‘historical global rate’ and ‘historical variable rate’ models. These historical models allow (but do not enforce) slow rates of colonisation leading to ‘over-dispersed’ assemblages that bear the signal of past allopatric speciation events, that is, close relatives co-occur less than expected if species presence/absence were randomly permuted. Failure to reject these historical models would therefore indicate that the slowdown in the empirical LTT can be explained as legacy of allopatric speciation and constant rates of colonisation (and local extinction) without needing to invoke a decline in colonisation rate over time.

This analysis tested whether slowdowns in community phylogenies patterns previously attributed to ecological limits could also be explained by a model accounting for speciation history but in which ecological limits are not explicitly assumed.

Cusimano, N. & Renner, S.S. (2010). Slowdowns in diversification rates from real phylogenies may not be real. *Systematic Biology*, 59, 458–464.

Etienne, R.S. & Haegeman, B. (2012). A conceptual and statistical framework for adaptive radiations with a key role for diversity dependence. *American Naturalist*, 180.

Etienne, R.S., Haegeman, B., Stadler, T., Aze, T., Pearson, P.N., Purvis, A., *et al.* (2012). Diversity-dependence brings molecular phylogenies closer to agreement with the fossil record. *Proceedings. Biological sciences / The Royal Society*, 279, 1300–9.

Mrkvička, T., Myllymäki, M. & Hahn, U. (2017). Multiple Monte Carlo testing, with applications in spatial point processes. *Stat Comput*, 27, 1239–1255.

Murrell, D.J. (2018). A global envelope test to detect non‐random bursts of trait evolution. *Methods Ecol Evol*, 9, 1739–1748.

Pigot, A.L. & Etienne, R.S. (2015). A new dynamic null model for phylogenetic community structure. *Ecology Letters*, 18, 153–163.

Price, T.D., Hooper, D.M., Buchanan, C.D., Johansson, U.S., Tietze, D.T., Alström, P., *et al.* (2014). Niche filling slows the diversification of Himalayan songbirds. *Nature*, 509, 222–225.

Weir, J.T. (2006). Divergent Timing and Patterns of Species Accumulation in Lowland and Highland Neotropical Birds. *Evolution*, 60, 842.

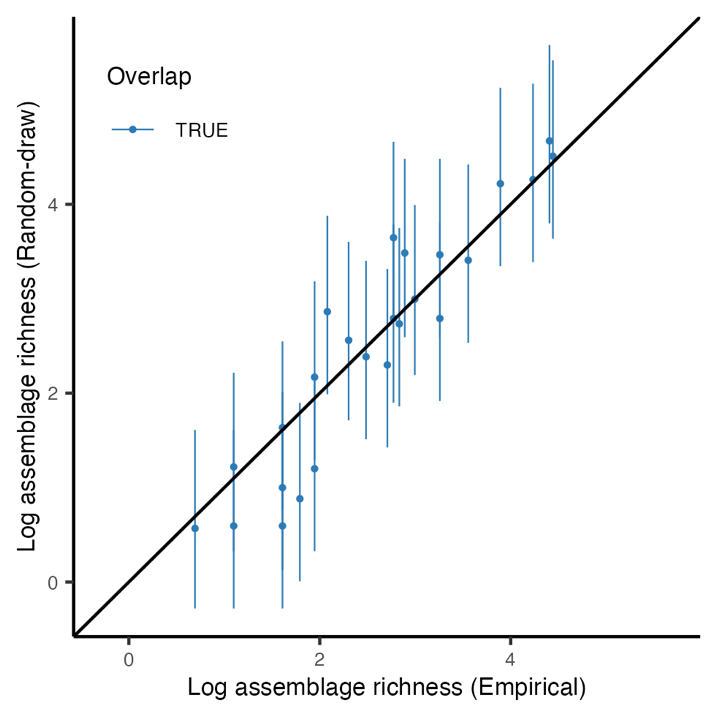

**Figure S1** Predicted versus maximum empirical sympatric diversity across the families at the 2600 km^2^ scale. The figure illustrates the strong positive link between assemblage and clade diversity. The predicted assemblage diversity results from a random-draw model in which the empirical proportions of maximum co-occurring diversity of each of the 25 families are randomly redistributed across the 25 families. This process is repeated a thousand times, and assemblage diversity is then calculated by multiplying the randomised proportions with the total species richness of the family. 95% Confidence intervals are calculated from the 1,000 replicates. The random-draw null model captures 25 out of 25 families.

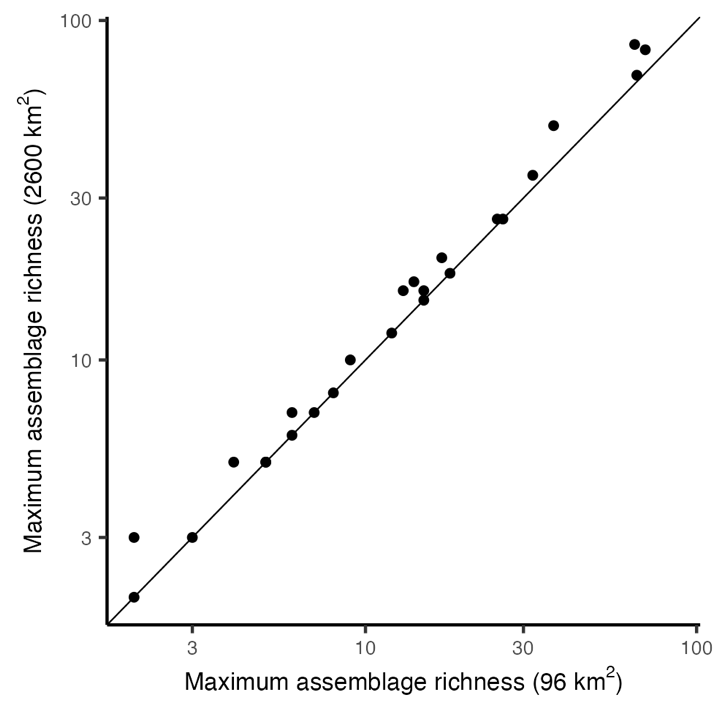

**Figure S2** Maximum assemblage diversity compared between grid cell sizes. The species richness of the grid cells with the maximum richness for all clades is compared to the richness for our two grid cell sizes (96 versus 2600 km^2^). The line through the origin indicates where species richness across scales is equal. The plot shows that richness across scales is marginally different for most families.

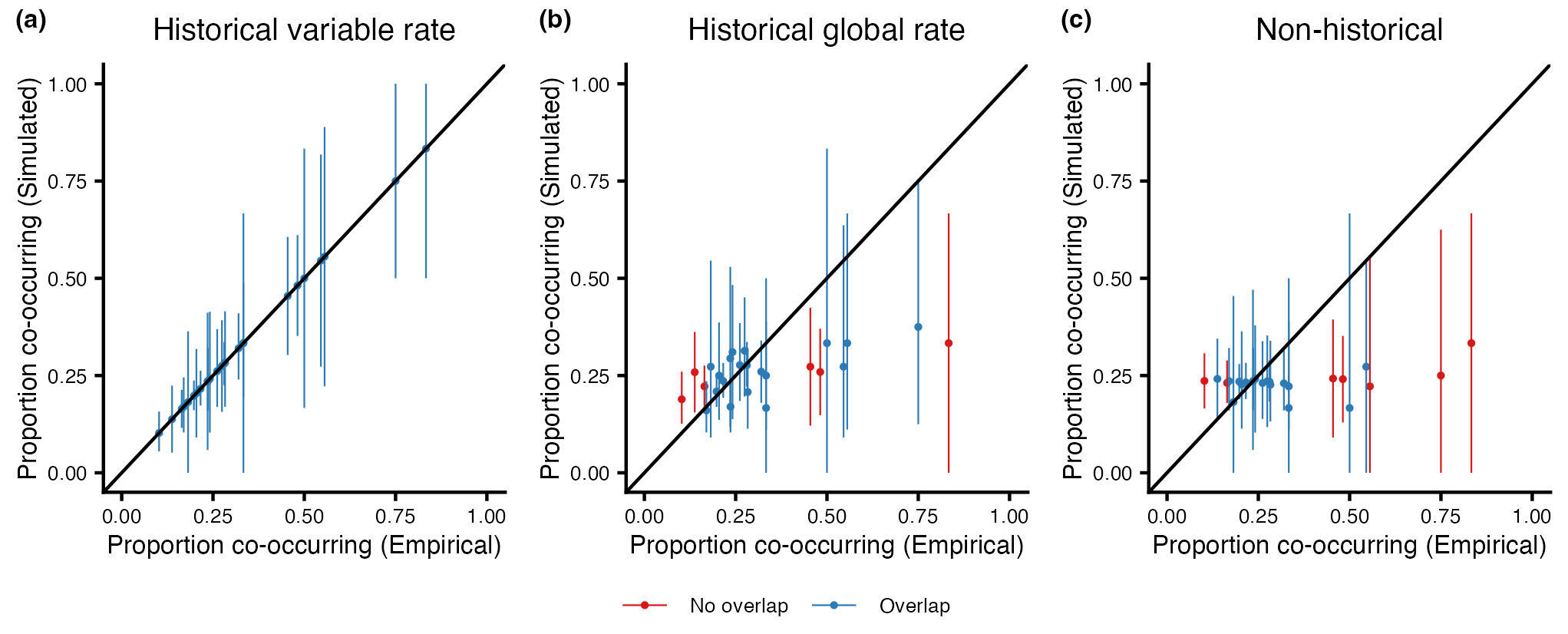

**Figure S3** Observed and predicted proportional richness of passerine clades at the 96 km^2^ scale. This figure is the equivalent of Figure 2, but at the 96 km^2^ instead of the 2600 km^2^ scale. For respectively (a,b and c) the predicted proportional assemblage diversity according to the historical variable rate, the historical global rate, and the non-historical models is plotted against the empirical proportions. Bars represent the 95% confidence intervals of the proportional richness of the community recovered under 2500 simulations. Colours indicate if empirical proportions fall within (blue) or outside (red) of the confidence intervals. For the non-historical model 25 out of 25 empirical proportion fall within the 95% confidence interval, whereas 23 out of 25 fall within the confidence intervals of the historical variable rate model and 19 out of 25 for the historical global rate model.

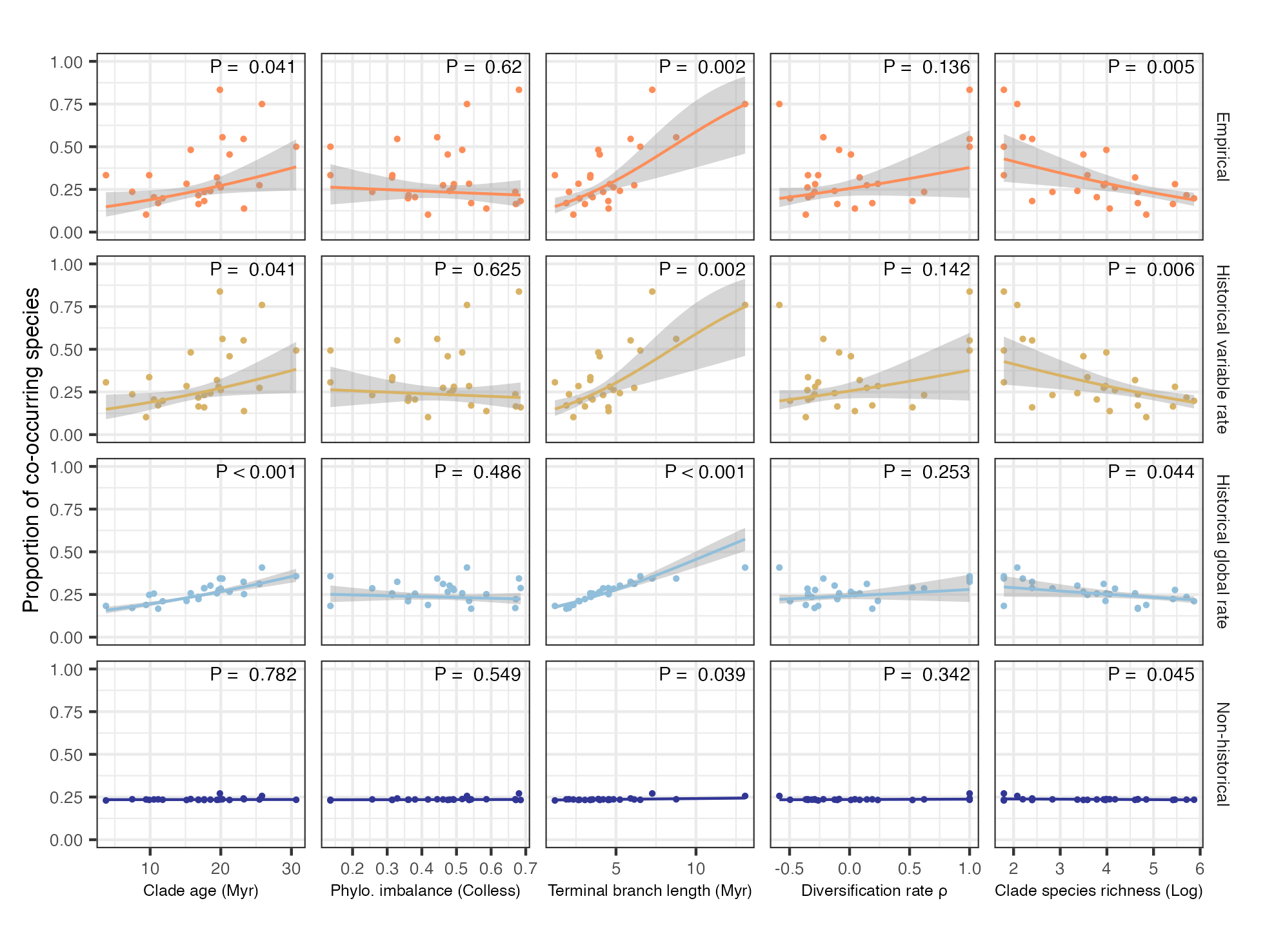

**Figure S4** Relationship between evolutionary history and proportional assemblage diversity at the 96 km^2^ scale. This figure is the equivalent of Figure 3, but at the 96 km^2^ instead of the 2600 km^2^ scale. Colours and rows denote the empirical relationships for all clades (orange) and relationships simulated under the historical variable rate (yellow), historical global rate (blue), and the non-historical (dark blue) models. For the simulation scenarios the mean proportion across 2500 simulations are plotted. Columns represent the five metrics representing evolutionary history: The crown age of the clade, phylogenetic imbalance (Colless’ index), mean branch length of the extant species, ρ a measure of temporal change in diversification rates through time where positive (negative) values indicate an increase (decrease), and log-transformed clade species richness. Shaded areas represent 95% confidence intervals.

**Table S1.** Parameter estimates for the historical global and variable rate models. Parameter estimates are given for the entire dataset and family-level phylogenies when conditioning on communities of at least 1 species (cond = 1) and when no conditioning is applied (cond = 0). 95% confidence intervals were calculated via parametric bootstrapping across 100 simulations.

| Data | N | Local N | Mu (cond = 0) | Mu  (cond = 1) (95%CI) | | Gamma  (cond = 0) | Gamma  (cond = 1) (95%CI) | |
| --- | --- | --- | --- | --- | --- | --- | --- | --- |
| Global | - | - | 0.175 | 0.178 | (0.082:0.302) | 0.12 | 0.12 | (0.093:0.17) |
| Calyptomenidae | 6 | 3 | 2.037 | 103.522 | (0:70.713) | 2.037 | 100.006 | (0.025:68.305) |
| Conopophagidae | 11 | 3 | 174.866 | 187.064 | (0:10.74) | 65.574 | 66.845 | (0.019:6.068) |
| Cotingidae | 65 | 20 | 0.008 | 0.008 | (0:1.008) | 0.056 | 0.056 | (0.037:0.516) |
| Dendrocolaptidae | 54 | 26 | 0 | 0 | (0:0.309) | 0.137 | 0.137 | (0.103:0.353) |
| Eurylaimidae | 8 | 6 | 65.46 | 1.934 | (0:5.905) | 196.356 | 5.803 | (0.042:13.059) |
| Formicariidae | 11 | 7 | 0 | 0 | (0:10.191) | 0.133 | 0.133 | (0.069:14.176) |
| Furnariidae | 225 | 49 | 0.635 | 0.635 | (0.099:82.898) | 0.204 | 0.204 | (0.079:26.68) |
| Grallariidae | 51 | 17 | 0.185 | 0.185 | (0:16.464) | 0.123 | 0.123 | (0.037:11.062) |
| Onychorhynchidae | 6 | 5 | 0 | 0 | (0:1.154) | 0.314 | 0.314 | (0.086:5.768) |
| Pipridae | 53 | 16 | 0.836 | 0.836 | (0:19.295) | 0.405 | 0.405 | (0.089:8.711) |
| Pipromorphidae | 100 | 35 | 0.033 | 0.033 | (0:6.509) | 0.097 | 0.097 | (0.065:2.551) |
| Pittidae | 29 | 7 | 0.191 | 0.191 | (0:19.245) | 0.086 | 0.086 | (0.02:8.604) |
| Platyrinchidae | 9 | 5 | 0.042 | 0.042 | (0:5.544) | 0.12 | 0.12 | (0.032:10.594) |
| Rhinocryptidae | 58 | 8 | 0.068 | 0.068 | (0:17.456) | 0.035 | 0.035 | (0.014:3.137) |
| Scleruridae | 17 | 5 | 0 | 0 | (0:12.627) | 0.052 | 0.052 | (0.012:7.352) |
| Thamnophilidae | 235 | 69 | 0.422 | 0.422 | (0.097:20.798) | 0.2 | 0.2 | (0.08:9.368) |
| Tityridae | 33 | 15 | 0.128 | 0.128 | (0:22.244) | 0.204 | 0.204 | (0.082:17.444) |
| Tyrannidae | 301 | 85 | 0.128 | 0.128 | (0:0.435) | 0.112 | 0.112 | (0.066:0.209) |
| Calcariidae | 6 | 2 | 0 | 0 | (0:804.035) | 0.156 | 0.138 | (0.138:329.985) |
| Cardinalidae | 44 | 10 | 0 | 0 | (0:34.739) | 0.052 | 0.052 | (0.028:11.51) |
| Emberizidae | 36 | 12 | 51.255 | 56.861 | (0:95.067) | 25.632 | 28.429 | (0.061:60.451) |
| Icteridae | 106 | 18 | 8.911 | 8.911 | (0.002:237.296) | 1.846 | 1.846 | (0.063:55.2) |
| Parulidae | 106 | 26 | 0 | 0 | (0:3.218) | 0.106 | 0.106 | (0.076:0.672) |
| Passerellidae | 127 | 16 | 0.545 | 0.545 | (0:673.994) | 0.109 | 0.109 | (0.031:111.302) |
| Thraupidae | 354 | 82 | 0.302 | 0.302 | (0.062:1.407) | 0.149 | 0.149 | (0.07:0.474) |

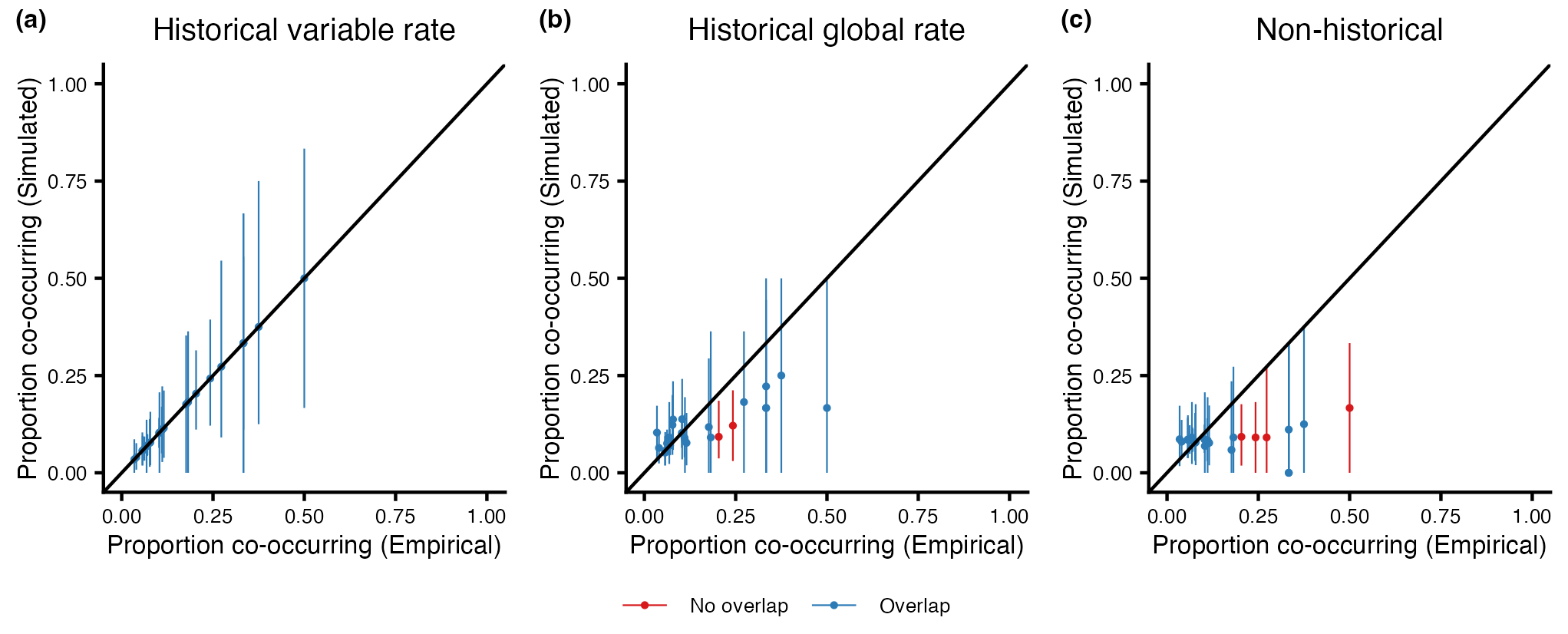

**Figure S5** Observed relative mean and predicted proportional richness and diversification rate of passerine clades. The figure is equivalent to Figure 2, but for each clade an assemblage of mean richness is used instead of maximum richness. For respectively (a, b and c) the predicted proportional assemblage diversity according to the historical variable rate, the historical global rate, and non-historical models are plotted against the empirical proportions. The line through the origin shows where the empirical equals the simulated proportion. Bars represent the 95% confidence intervals of the proportional richness of the assemblages recovered under 2500 simulations. Colours indicate if empirical proportions fall within (blue) or outside (red) of the confidence intervals. For respectively the historical variable rate (AIC = 1204.018), the historical global (AIC = 1162.246) rate, and the non-historical model (AIC = 1205.494) (a) 25 out of 25, (b) 23 out of 25, and (c) 21 out of 25 fall within the confidence intervals for sympatric diversity.

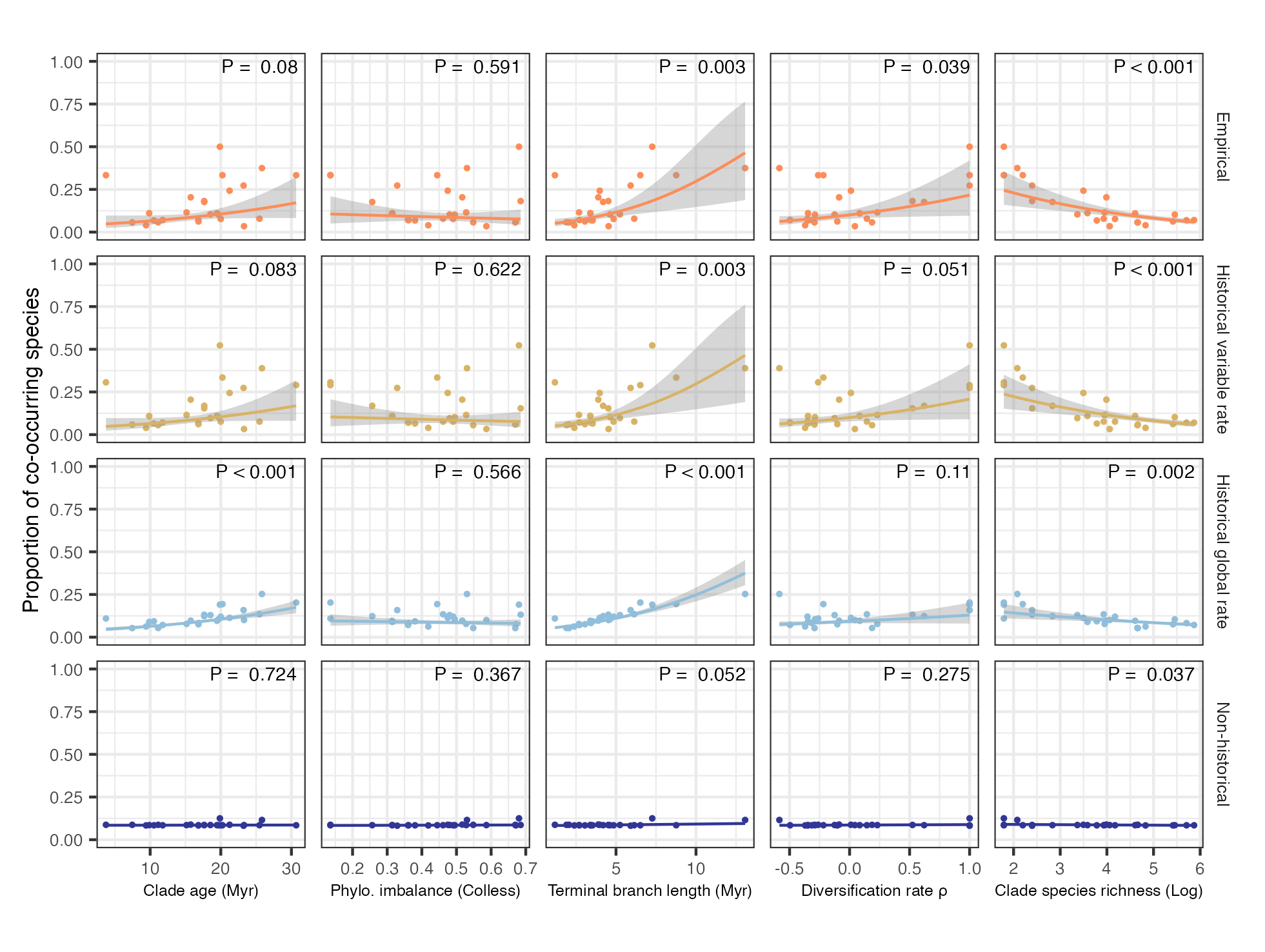

**Figure S6** Relationship between evolutionary history and proportional mean assemblage diversity. The figure is equivalent to Figure 3, but for each clade an assemblage of mean richness is used instead of maximum richness. Colours and rows denote the empirical relationships for all clades (orange) and relationships simulated under the historical variable rate (yellow), historical global rate (blue), and the non-historical (dark blue) models. For the simulation scenarios the mean proportion across 2500 simulations are plotted. Columns represent the five metrics representing evolutionary history: the crown age of the clade, phylogenetic imbalance (Colless’ index), mean branch length of the extant species, ρ which is a measure of temporal change in diversification rates through time where positive (negative) values indicate an increase (decrease), and log-transformed clade species richness. Fitted Generalised Linear Models are shown, and shaded areas represent 95% confidence intervals.

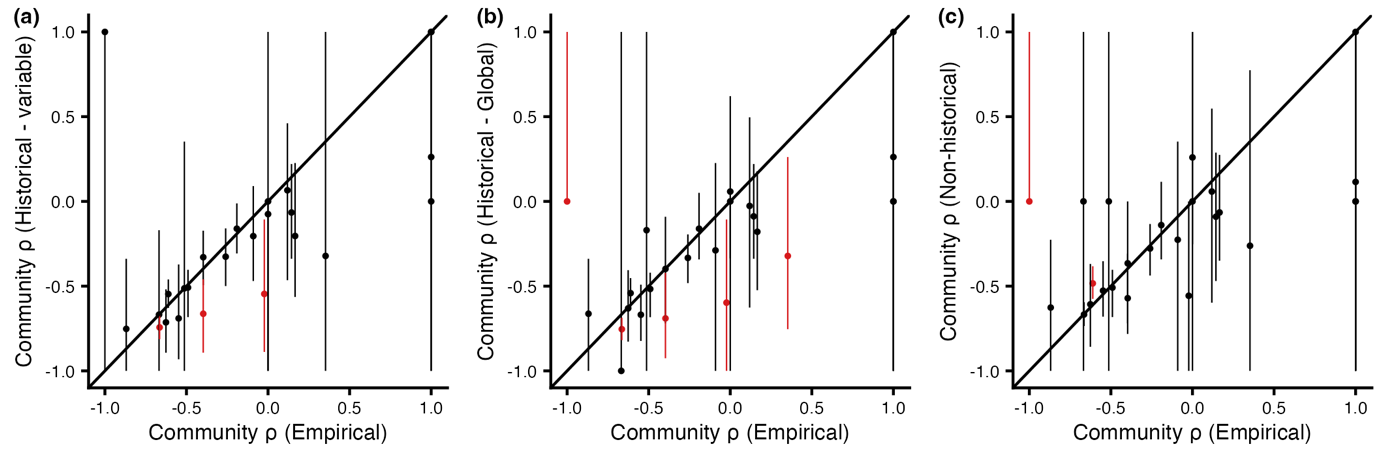

**Figure S7** Comparisons of simulated and empirical diversification rate patterns. Results represent the same set of simulations as shown in Figure 2 at the 2600 km^2^ scale. On the *x*-axis of each plot the empirical ρ is shown, calculated by pruning the family-level phylogenies of any species that do not occur in the assemblage of highest sympatric species richness. Panels show the estimated ρ across 2500 phylogenies pruned by removing the species absent from the simulated assemblages for respectively the (a) the historical variable rate model (22/25), (b) the historical global rate model (20/25), and (c) the non-historical model (23/25). Negative values of ρ indicate slowdowns in diversification rate through time, whereas positive values indicate increases.

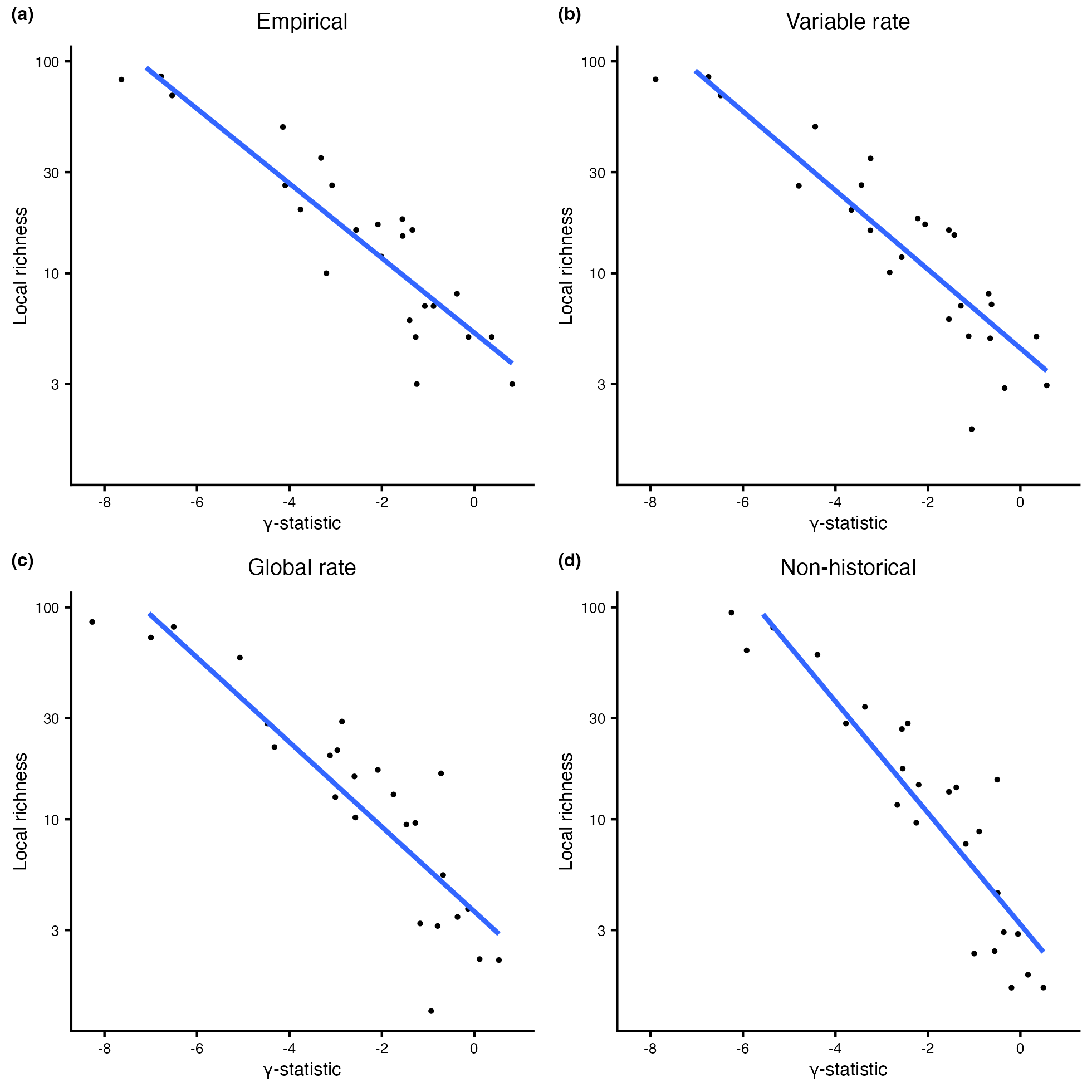

**Figure S8** The relationship between the γ-statistic and assemblage richness. Relationships are shown between the assemblage-level (2600 km^2^) γ -statistic, where negative values indicate slowdowns in diversification rate. To calculate γ, phylogenies were pruned to only include species that were present in the geographical cell with the highest species richness (*x*-axis) of the respective families for the empirical data, or the simulated presence for the simulation scenarios. The scenarios from left to right, top to bottom, represent (a) the empirical relationship, (b) historical variable rate model, (c) the historical global rate model, and (d) the non-historical model. Trends shown are significant linear regression fits.
